# Queen and king recognition in the subterranean termite, *Reticulitermes flavipes*: Evidence for royal recognition pheromones

**DOI:** 10.1101/496240

**Authors:** Colin F. Funaro, Coby Schal, Edward L. Vargo

## Abstract

Royal recognition is a central feature of insect societies, allowing them to maintain the reproductive division of labor and regulate colony demography. Queen recognition has been broadly demonstrated and queen recognition pheromones have been identified in social hymenopterans, and in one termite species. Here we describe behaviors that are elicited in workers and soldiers by neotenic queens and kings of the subterranean termite, *Reticulitermes flavipes*, and demonstrate the chemical basis for the behavior. Workers and soldiers readily perform a lateral or longitudinal shaking behavior upon antennal contact with queens and kings. When royal cuticular chemicals are transferred to live workers or inert glass dummies, they elicit antennation and shaking in a dose-dependent manner. The striking response to reproductives and their cuticular extracts suggests that royal-specific cuticular compounds act as recognition pheromones and that shaking behavior is a clear and measurable queen and king recognition response in this termite species.

## Introduction

Social insects rely on chemical communication to function effectively; within the colony, pheromones mediate foraging, aggregation, defense, reproduction, and other essential processes [1,2]. Recognizing reproductive castes (queens in social hymenopterans, queens and kings in termites) is especially important to preserve the royal-worker division of labor and to ensure proper care for these high-value individuals. Royal pheromones and recognition behaviors have been well studied in ants, bees, and wasps, but have received little attention in termites. Pheromones largely mediate and guide the behavior and physiology of sterile castes in social hymenopteran colonies, with some notable exceptions that include visual signals and tactile/physical interactions [3–5]. Pheromones in general and more specifically royal (usually queen) pheromones are generally classified into those that elicit immediate behaviors (releaser pheromones) and those that induce long-term physiological changes in sterile worker castes (primer pheromones). Identifying these compounds and elucidating their effects and glandular origins have received increasing attention in social insect biology.

Distantly related to the social Hymenoptera, termites share life history traits and ecologically important roles with ants, bees, and wasps. Termites tend to exhibit a more flexible developmental pathway than their hymenopteran counterparts, as most individuals in colonies of many lower termites, including subterranean termites (Blattodea: Isoptera: Rhinotermitidae), retain the ability to develop gonads and molt into functional worker-derived reproductives called neotenics [6,7]. Neotenics can also develop from nymphs that normally develop into winged primary reproductives. Active queens and kings inhibit reproductive development in other colony members and most likely use chemical signals to do so. Additionally, because reproductively active males (kings) stay within the nest, termites appear to employ both queen- and king-specific pheromones to preserve the reproductive division of labor in each sex and elicit care from workers [8–10].

The first termite queen primer pheromone was identified in the Japanese subterranean termite (*Reticulitermes speratus*) as a blend of two highly volatile compounds—2-methyl-1-butanol and n-buty1-n-butyrate—which inhibit the reproductive differentiation of female workers and nymphs into supplementary reproductives [10]. Although reproductive-specific volatile compounds and long-chain hydrocarbons have been found in *Nasutitermes takasagoensis* and *Zootermopsis nevadensis*, respectively, their functions have not been evaluated [11,12]. Thus, only one releaser pheromone involved in royal recognition has recently been described in termites [13]. It is possible that the search for these compounds has been impeded by the rarity and fragility of termite reproductives, the paucity of termite researchers, or the lack of robust bioassays to measure the physiological or behavioral effects of presumptive queen and king pheromones.

Foundational work on queen-recognition in bees and fire ants involves ketones, esters, alcohols, and fatty acids [14,15]. However, queens from a number of other social hymenopteran species possess unique cuticular hydrocarbons (CHCs) that correlate with ovary activation and often elicit queen recognition [16–18]. CHCs are the dominant class of most insects’ cuticular lipid layer and help to prevent desiccation and act as a barrier against pathogens. Also known to be important cues in nestmate and interspecific recognition in both solitary and social species, CHCs are highly variable and responsive to different physiological and environmental inputs [19]. CHCs are thus hypothesized to have been co-opted over the course of evolution as reliable signals for mate recognition and fertility because of the relationship between their composition and the insect’s physiology and metabolism [17,20].

Cuticular hydrocarbons also show promise as termite royal recognition pheromones. They have been found to be involved in caste- and species-recognition, especially within the subterranean termites [21–25]. The literature remains divided on the role of CHCs in recognition and aggression, but it is generally held that CHC blends are important in mediating behavior and agonistic interactions in certain termite species [26,27]. Indeed, the only royal recognition pheromone, recently identified in the subterranean termite *Reticulitermes flavipes*, is a CHC [13].

Pivotal to the identification of royal-recognition pheromones are behavioral assays that can be used to test worker responses to queens and kings. Yet, queen and king recognition behaviors have not been well described in termites, and there is no clear retinue response around reproductive individuals, as commonly observed in the Hymenoptera [15,28–30]. Aggressive intracolony interactions establishing reproductive dominance have been described and are loosely related to queen recognition, but it is unclear whether a chemical signal is involved [31,32]. Behaviors such as head-butting, tremulations, jerking, and oscillatory behavior have been described in several termite species, but never in relation to reproductive individuals [33–37].

Upon observing a distinct oscillatory/shaking behavior in workers of *R. flavipes* in close proximity to primary and neotenic queens and kings, we hypothesized that this might represent a royal recognition response. To complement a growing literature describing queen fertility and recognition signals in social Hymenoptera and to understand the mechanisms underlying royal recognition in termites, we investigated behavioral responses of workers to queens and kings of *R. flavipes* and the chemical signals that elicit them.

## Methods and Materials

### Termite collection

Colonies of *Reticulitermes flavipes* were collected as needed in Raleigh, North Carolina, from three wooded locations (Carl Alwin Schenck Memorial Forest, 35.48 N, 78.43 W, under the authority of Elizabeth Snider; Historic Yates Mill County Park, 35.43 N, 78.41 W, under the authority of Timothy Lisk; and Lake Johnson Park, 35.45 N, 78.42 W) between 2010 and 2015. Lake Johnson Park is managed by the city of Raleigh and requires no specific permissions for termite collection and none of our collections involved endangered or protected species at any collecting site. Although colonies could be maintained for long periods of time and successfully reproduced in the lab, they tended to slowly lose their vitality and needed to be replaced. Additionally, primary queens and kings were rarely found, and colonies were most likely not collected in their entirety because of the many spatially separated chambers typical of *R. flavipes* colonies. For each assay, specific colonies and their collection site are noted in the methods below. All termite colonies were maintained in laboratory conditions for ~6 months before use, with the exception that the initial behavioral assays used colonies from Schenck Forest maintained for ~11 months in the lab. Whole tree limbs or logs with termites were split into smaller pieces and set out in shallow pans to dry. Using either plastic container lids with moist paper towels underneath or ~10 cm PVC pipes containing coils of moistened corrugated cardboard, the termites passively moved out of the drying wood and into the moist substrate. Fully extracted colonies were kept either in clear plastic boxes lined with moist sand and pine shims for food or in 9-cm petri dishes with an autoclaved lab substrate consisting of 70% sawdust and 30% α-cellulose. Colonies were maintained in opaque plastic containers in a ~24°C incubator under 14:10 L:D cycle with lights-on at 0600.

### Production of secondary reproductives

Shaking behavior in response to royal castes was initially observed in primary, colony founding queens and kings. Because these individuals are difficult to find and *R. flavipes* readily generates replacement, or neotenic, reproductives, we used only neotenic kings and queens in our assays. To produce these individuals, colony fragments of ~2,000 to 5,000 individuals were subdivided into 5 cm petri dishes without reproductives. Newly emerged neotenics typically appeared within 2–3 weeks and were removed to prevent inhibition of queen and king differentiation in the neotenic-generating dishes. Newly-emerged neotenics were then held in 9-cm dishes containing ~500 workers and 20–50 soldiers until used in experiments. The majority of neotenics were ergatoid (and thus apterous), although on rare occasions we found nymphoid neotenics present in the colony fragments, which were identified by their wingbuds. Additionally, neotenic kings typically differentiated one at a time, while multiple female neotenics would differentiate simultaneously from one dish, which limited the scope of our king based experiments.

### General bioassay procedure

Unless noted otherwise, termites were divided into dishes of 30 workers and two soldiers, allowed to acclimate for at least 7 days and observed once. To lessen the statistical effects of repeated measures and to more efficiently use the available termites, we assayed 10 replicate dishes per treatment and then returned the termites into a larger colony. Then we re-distributed the termites from the larger colony back into new replicate dishes. This allowed us to use the termite workers several times by randomizing individuals across replicates and treatments.

We assessed the differential responses of worker and soldier termites to neotenic queens and kings, workers, and soldiers. Behaviors that were readily observed and quantified included total time spent moving by the focal termite and the total number of allogrooming sessions, shaking responses, and antennations by both the focal termite (active) and by resident workers and soldiers that interacted with it (reactive). Of these, reactive shaking by resident termites in response to the introduced focal termite emerged as the most discriminating and showed a clear and significant difference between reproductive and non-reproductive individuals (below). Shaking behavior was defined as repetitive lateral oscillatory movements. All active shaking behaviors performed by the focal termite were recorded, while reactive shaking behaviors in resident termites were only recorded when they were within approximately 1 mm of the focal individual. Whereas shaking responses by the resident termites (reactive shaking) significantly discriminated royal and non-royal castes, shaking responses by the focal termite (active shaking) varied across castes, but failed to discriminate workers and soldiers from queens (Fig 1A,B). Therefore, only shaking responses by resident termites in response to the introduced focal termite were used in subsequent assays. Antennation responses of resident termites were also informative, though to a lesser extent than shaking responses. Nevertheless, in all subsequent results we report on both reactive shaking and reactive antennation responses. We defined antennation as placing both antennae on another individual and any continuous contact was counted as one session. Termites had to move more than ~3 mm away from the focal termite before we recorded a second antennation session.

**Fig 1.**
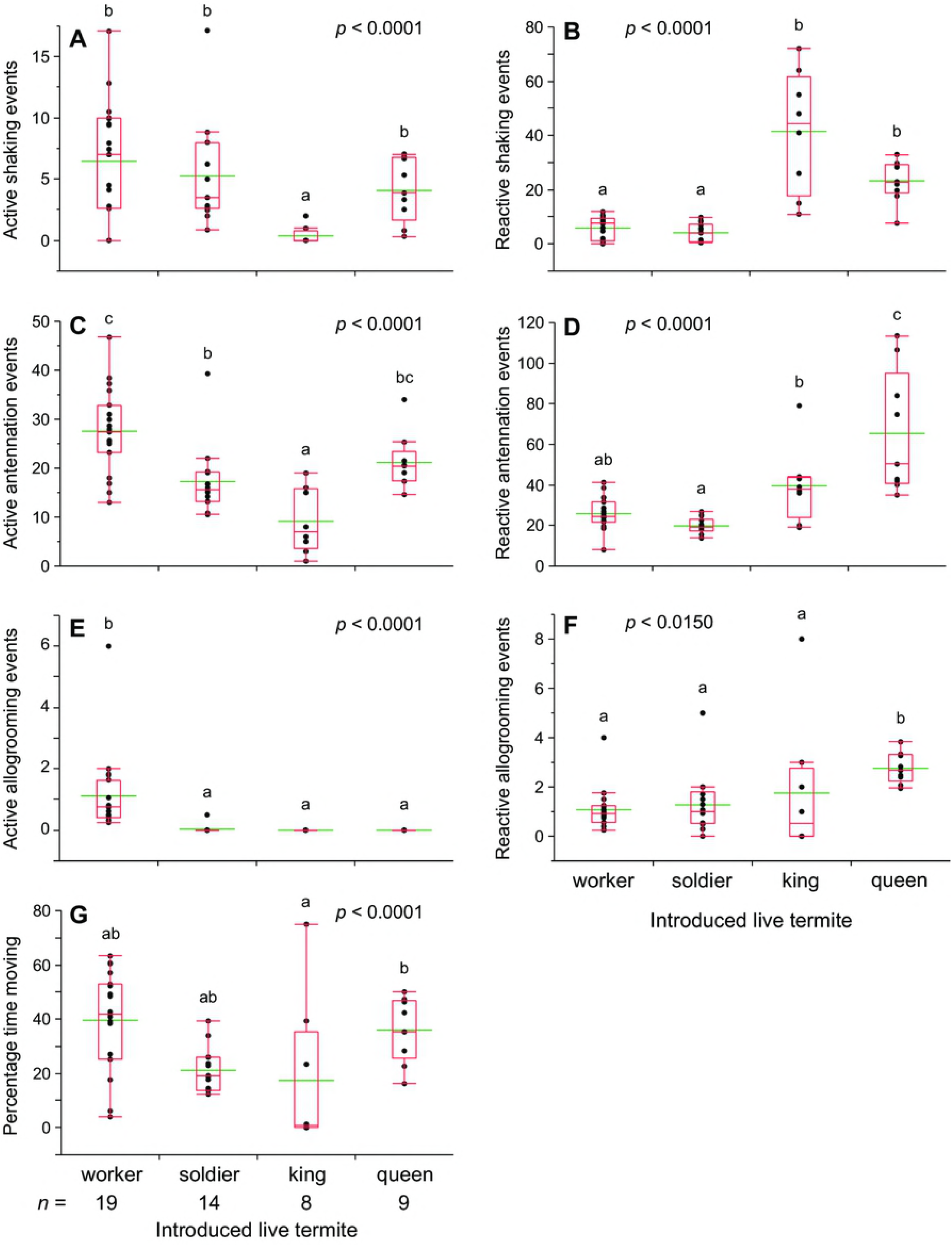
Termite behavioral responses to a worker, soldier, neotenic king, and neotenic queen. Behavioral responses were measured in 10 min assays. Queens and kings in all assays were neotenic (secondary) reproductives generated within the lab. Each assay dish consisted of 30 workers, 2 soldiers, and an introduced live focal termite, and assays were conducted under ambient light conditions during the photophase. For all treatments the number of replicate assays is indicated under the axis for each caste in (**G**). Letters indicate significantly different values using one-way ANOVA (*p* < 0.05) and Tukey’s HSD. In the box plots, the horizontal line within the box represents the median value, the box represents the 25th to 75th quantiles, and the wider green line represents the mean.

### Royal recognition bioassay

All termites in these assays were from a single colony collected in Schenck Forest in 2010. Queens and kings used in recognition assays were neotenics, as primary reproductives were too rare to provide effective replication. One hundred workers were placed into a 5-cm petri dish with moist unwoven paper towels or filter paper. Two soldiers were added to each dish to discourage soldier differentiation during the experiment. To differentiate the introduced (focal) worker from resident workers, the introduced worker was dyed blue by feeding on substrate impregnated with 0.1% Nile Blue. Workers turned blue in one to two weeks of consuming dyed diet and appeared to show no decrease in activity or survivorship, as previously described [38]. A single dyed worker was introduced to each assay dish at least 24 h before the assay. Assays consisted of removing the lid of the petri dish, wiping off any condensation to improve visibility, waiting at least 2 min to rest the termites, and observing the dyed focal worker for 10 min. Measured parameters included total time spent moving by the focal worker, total number of distinct allogrooming sessions, shaking response, as measured by the number of shaking events during the assay, and antennation response, as measured by the number of antennation events during the assay. All behaviors (allogrooming, shaking, and antennation) were measured in both the focal workers of the assay and the resident workers that interacted with it, leading to observed behaviors being divided into either active (actions performed by the focal termite) or reactive (reactions elicited by the focal termite from others) categories.

### Behavioral assays in light and dark conditions

An experiment was conducted to examine the behavioral responses of workers to queens in the photophase and scotophase, and under light and dark conditions. Termites from a colony collected at Lake Johnson Park in 2015 were placed in a 5-cm petri dish (30 workers and 2 soldiers), observed once, and returned to the colony for re-use. A queen was introduced into each petri dish and observed either in a fully lit laboratory (~450 lux at the assay dishes) or within a dark box with only a red headlamp. Antennation and shaking elicited in resident termites were measured for 7 min.

### Foreign queen recognition bioassay

We performed assays similar to our royal recognition assays to test the responses of nestmate termites to unrelated individuals in dishes with 30 nestmate workers and two nestmate soldiers. Termites were collected in 2015 from colonies found at Lake Johnson Park and Schenck Forest, respectively, which are approximately 6.5 km apart. These assays were designed to assess the queen recognition activity of foreign queens versus native queens and to support observations that *R. flavipes* colonies show no observable aggression toward foreign queens. Petri dish lids were removed and condensation was wiped from the lid before one of four treatments was added: nestmate neotenic queen, nestmate worker, foreign neotenic queen, or foreign worker. Observations began immediately after a focal termite was introduced. Replicates were observed for 7 min and cumulative antennation and shaking elicited in resident termites were recorded each min. Introduced workers were dyed blue for tracking purposes. All treatments had 10 replicate petri dishes. Dishes were assayed once and then re-distributed into a larger colony. Because only five queens were available at the time of this assay, each queen was observed in two replicate assays 1 week apart.

### Transfer of cuticular compounds to live termites

To test whether the recognition behaviors elicited by queens were mediated by cuticular compounds and whether these compounds could be transferred to non-reproductive termites, we “perfumed” workers by tumbling queens with workers in various ratios. All termites used were collected in 2015 from a single colony from Lake Johnson Park in Raleigh. We included queen:worker ratios of 7:15, 1:1, 5:1, and 10:1 with a 40:15 worker:dyed worker negative control and a live tumbled queen as positive control. Each treatment (queen:worker ratio) and control was replicated 10 times. For the 7:15 queen:worker and 40:15 worker:worker experiments, 15 blue-dyed workers were tumbled in glass vials with either 7 queens or 40 workers, respectively. After perfuming, five blue-dyed workers were frozen for extraction and cuticular analysis and 10 were removed to a clean petri dish for use in the bioassay. For all other ratios (1:1, 5:1, and 10:1), a blue-dyed worker was tumbled alone with 1, 5, or 10 queens and then observed in the bioassay. Neotenic queens were derived from the native colony in all treatments except for the 10:1 queen:worker treatment, where three of the 10 queens were from a foreign colony. Cuticular compounds were transferred by placing live termites into 4 mL glass vials and gently rotating the vials to tumble them for 3 min. Efforts were made to maximize contact among termites. Queens were rested in the dark while running the assay and 5 tumbling sessions were performed each day to minimize stress in the queens. After tumbling, a single queen-“perfumed” dyed worker was added to each petri dish with 30 workers and two soldiers and immediately assayed for 7 min, recording cumulative antennation and shaking responses elicited in resident termites. As above, termites were assayed once and then re-distributed into a larger colony.

### Transfer of cuticular compounds to glass dummies

We designed an experiment similar to the previous queen compound transfer bioassay to test whether queen cuticular compounds could be extracted in hexane and effectively transferred to glass dummies. We melted Pasteur pipette tips into roughly the length and diameter of a neotenic queen (~2 mm x ~6 mm). Neotenic queens, neotenic kings, and workers were extracted in hexane (200 μL/individual) for 2 min with gentle mixing. Hexane was transferred to new vials and evaporated under a gentle stream of high purity nitrogen. Final concentrations of 0.1, 0.3, 1, and 3 queen- or king-equivalents (QE or KE) per 20 μL were created from the initial extract for a dose-response study of royal compounds. Worker controls were tested using 6 worker equivalents (WE) because worker body mass and CHC mass (CF, unpublished results) were approximately half those of queens and this would be equivalent to our highest concentration in royals. The bioassays tested one dummy per petri dish with two colonies (n = 10 dishes) for each queen treatment and controls and n = 5 dishes per colony for all king treatments due to limited availability of kings. First, glass dummies were rinsed in hexane and allowed to dry before applying 20 μL of extract onto each in a glass petri dish. Hexane was allowed to evaporate from treated dummies for 5 min before introduction into assay dishes. Observations began 2 min after introducing the dummy to allow the termites to settle. We measured antennation, shaking responses, and presence/absence of aggression towards the dummies for 5 min. Aggression was defined as a repetitive lunging motion toward the dummy. In these assays, each group of termites in a dish was observed once per treatment, but then observed again in five other treatments, with a rest period of at least 24 h between assays. This experiment was performed with termites collected in 2013 from two colonies (one from Schenck Forest and one from Yates Mill Park) and the data from both colonies were combined when no differences were found between the colonies.

### Statistical Methods

Comparisons made across treatments were analyzed with ANOVA with a post-hoc Tukey’s honest significant difference test. All assay count data was square root transformed. Aggression data were analyzed using a chi-square test, as the behavior was recorded as either present or absent. All statistical tests were run in JMP (JMP®, Version 12. SAS Institute Inc., Cary, NC, 1989-2007). Raw data for all experiments is available in the supporting information (S1 Dataset).

## Results

### Royal recognition bioassays and behaviors elicited by different castes

We assessed the differential responses of worker and soldier termites to neotenic queens and kings, workers, and soldiers (Fig 1). Shaking behavior occurred ~5–8-fold more in response to a neotenic queen or king than to a worker or soldier (Fig 1B). Differences in antennation were less pronounced, showing a ~2–3-fold increase in response to reproductives (Fig 1D). Though not pursued in other assays, allogrooming and movement rates both showed patterns across caste. Grooming by the focal termite (active grooming) was almost exclusively performed by workers and the introduced focal workers groomed resident termites significantly more than other castes (Fig 1E). Queens elicited significantly more allogrooming (reactive allogrooming) than workers, soldiers, or kings (Fig 1F). Workers moved around the assay dish significantly more than soldiers and kings, and both workers and queens spent ~2X more time moving in the assay dish than other castes (Fig 1G). While significant differences were found between castes for multiple behaviors, reactive shaking and antennation were the primary indicators of royal status.

### Behavioral assays in light and dark conditions

To optimize the behavioral assay we conducted observations in the photophase and scotophase and under light and dark conditions. There were no significant differences in the shaking (Fig 2A) or antennation responses (Fig 2B) to live neotenic queens between the photophase and scotophase. Observations under dark conditions in both photophase and scotophase, however, yielded significantly lower rates of shaking toward live neotenic queens than in a lit room. While there were no differences in antennation during the photophase, observations during the scotophase also showed significantly lower rates of antennation under dark conditions. Therefore, all subsequent assays were conducted under ambient light conditions during the photophase.

**Fig 2.**
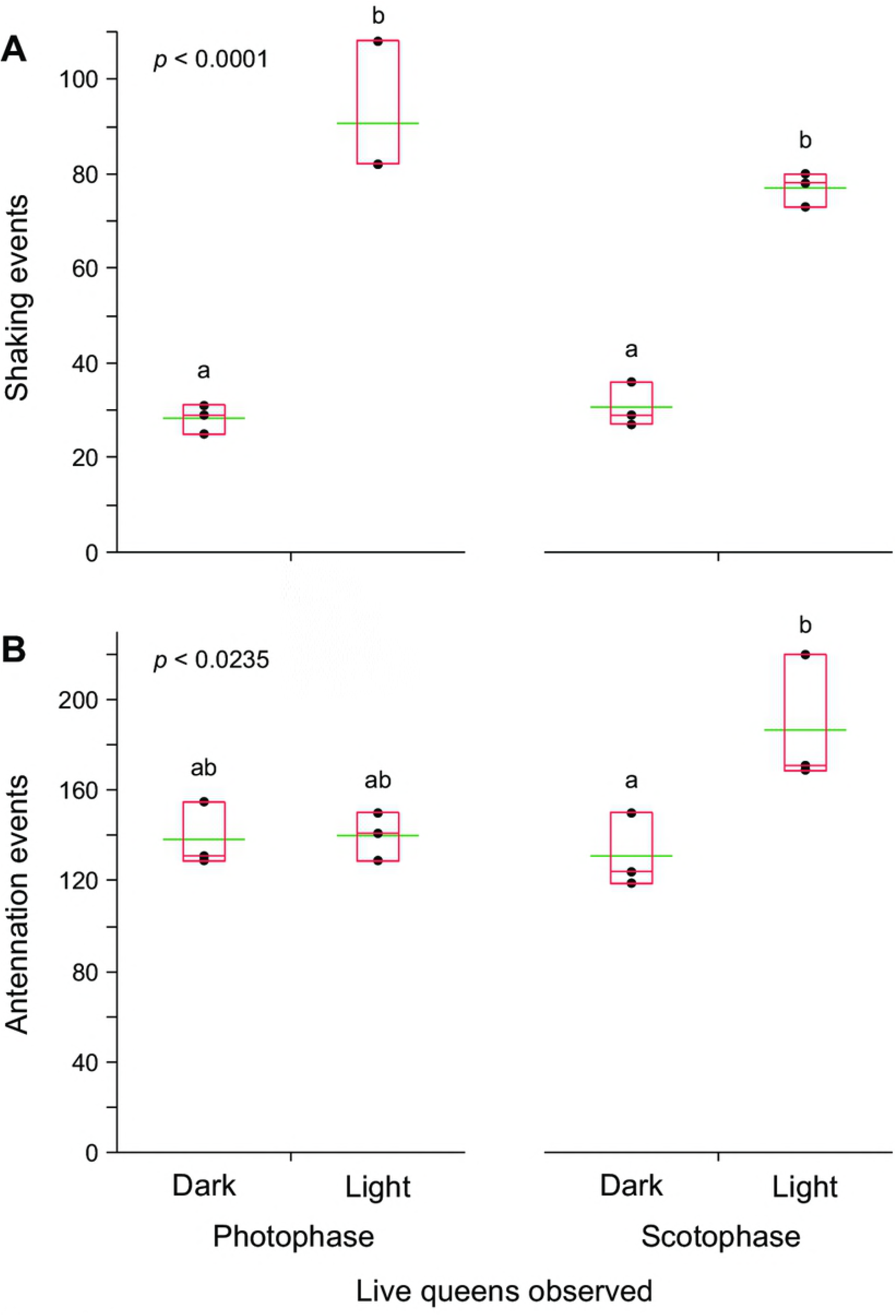
Response of termites to a live neotenic queen in light and dark conditions. Each assay dish consisted of 30 workers, 2 soldiers, and a live queen observed for 7 minutes. Termites were assayed in their photophase and scotophase and under light and dark conditions measuring both shaking (**A**) and antennation responses (**B**). The number of replicate assays was 3. Letters indicate significantly different values using one-way ANOVA (*p* < 0.05) and Tukey’s HSD. In the box plots, the horizontal line within the box represents the median value, the box represents the 25th to 75th quantiles, and the wider green line represents the mean.

### Foreign queen bioassay: workers respond similarly to native and foreign queens

In assays comparing responses to native and foreign workers and neotenic queens, workers and soldiers showed no overt aggression toward queens or workers introduced to dishes during the assay (CF, personal observations). However, both nestmate and foreign neotenic queens elicited strong shaking responses that were about four times higher than those elicited by nestmate and foreign workers (Fig 3A). Likewise, more antennation responses were elicited by nestmate neotenic queens than by nestmate workers, and foreign queens elicited more antennation responses than foreign workers (Fig 3B). Therefore, in some subsequent assays, foreign neotenic queens, which elicited strong responses similar to nestmate queens, were used, as noted, when nestmate queens were not available.

**Fig 3.**
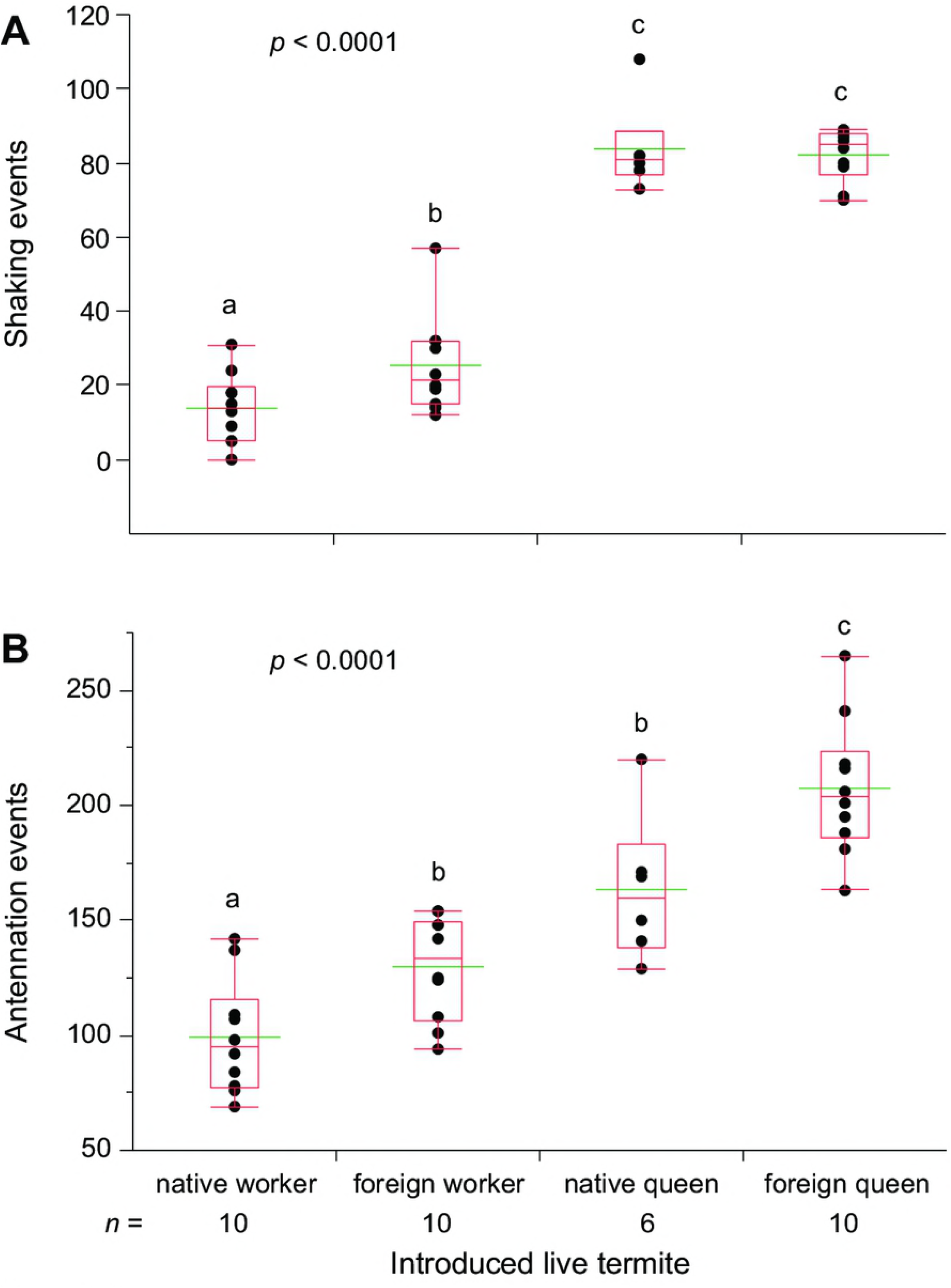
Termite behavioral responses to native and foreign neotenic queen with foreign and native worker controls. Shaking (**A**) and antennation (**B**) were measured in 7 min assays. Each assay dish consisted of 30 workers, 2 soldiers, and an introduced live focal termite, and assays were conducted under ambient light conditions during the photophase. The number of replicate assays is indicated under the axis for each caste. Letters indicate significantly different values using one-way ANOVA (*p* < 0.05) and Tukey’s HSD. In the box plots, the horizontal line within the box represents the median value, the box represents the 25th to 75th quantiles, and the wider green line represents the mean.

### Transfer of cuticular compounds to live termites and glass dummies: queen and king extracts elicit royal recognition

To test whether queen recognition compounds could be transferred from the queen to workers, we tumbled workers with neotenic queens in glass vials to transfer queen-specific CHCs to workers, or “perfume” them with queen scent. As negative controls, we tumbled 40:15 undyed workers with blue-dyed “focal” workers (i.e., 2.7X) to account for the greater body mass of queens. Dose-response assays included 0.5X to 10X queen:worker ratios, and a tumbled live queen represented the positive control. Queen-coated workers elicited significantly more shaking responses (Fig 4A) and antennation (Fig 4B) than worker-perfumed control workers. Notably, there was a clear dose-response relationship between the queen:worker ratio per tumbled worker and shaking responses (Fig 4A). However, despite the effective transfer of queen compounds to workers, none of the queen-perfumed workers elicited a shaking response as strong as live neotenic queens. Kings were not tested in these assays due to the small number of them available.

**Fig 4.**
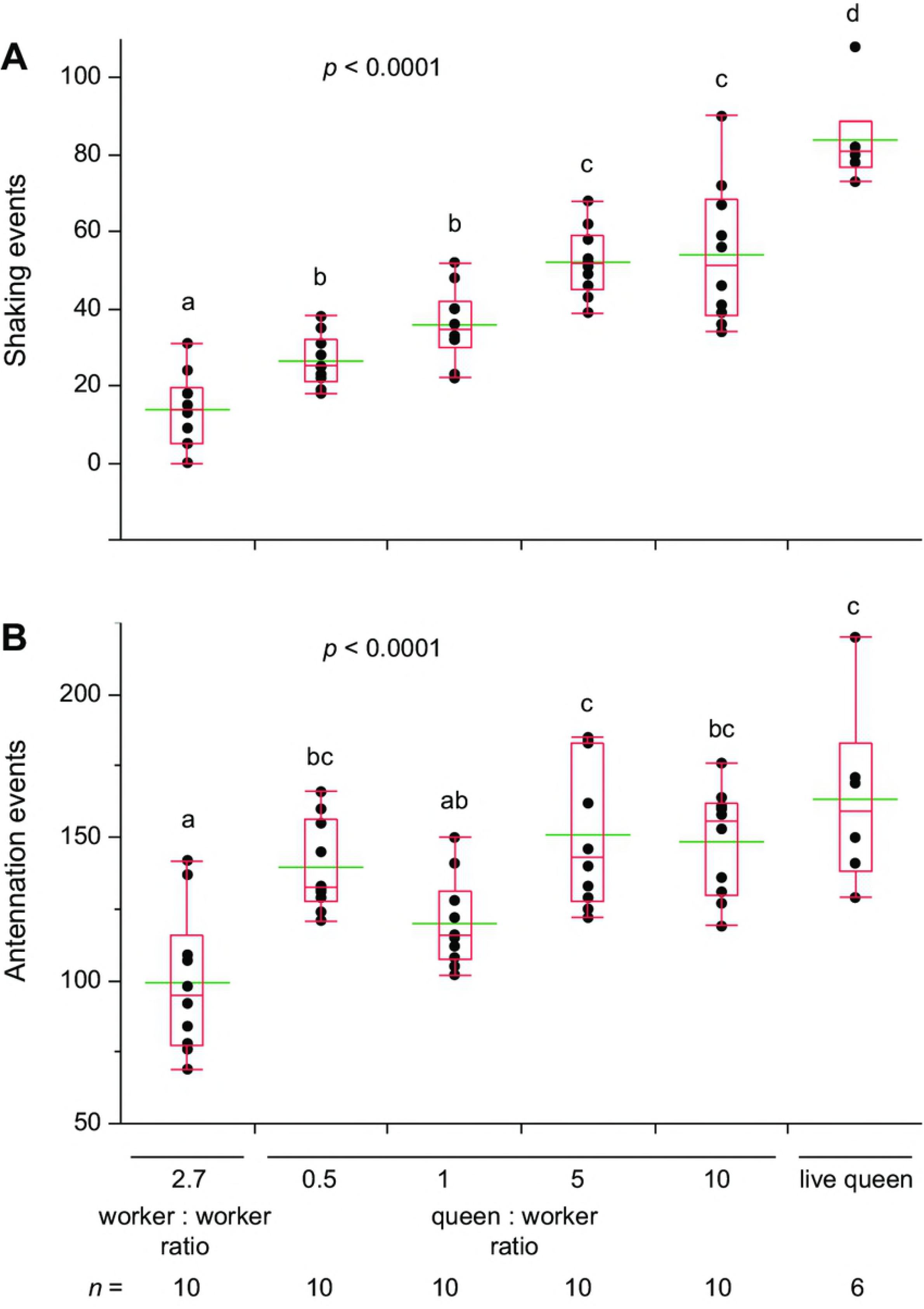
Termite responses to live workers coated with neotenic queen cuticular compounds. Queen recognition behavior was measured in 7 min assays via shaking (**A**) and antennation (**B**) responses. Workers were tumbled in clean glass vials with other workers at a ratio of 40:15 (2.7X) dyed workers: undyed workers or with queens at ratios of Queen: Worker of 7:15 (0.5X), 1:1 (1X), 5:1 (5X), and 10:1 (10X). A live neotenic queen was tumbled in a vial as a positive control. Each assay dish consisted of 30 workers, 2 soldiers, and a tumbled test individual, and assays were conducted under ambient light conditions during the photophase. For all treatments the number of replicate assays is indicated under its axis label. Letters indicate significantly different values using one-way ANOVA (*p* < 0.05) and Tukey’s HSD. In the box plots, the horizontal line within the box represents the median value, the box represents the 25th to 75th quantiles, and the wider green line represents the mean.

Finally, to control for the presence of non-chemical cues on workers that might facilitate the queen recognition responses, we transferred hexane extracts of workers and neotenic queens and kings to glass dummies, which were introduced into assay dishes. We used the extract of 6 workers (6 WE) as negative control and 0.1 to 3 neotenic queen- or king-equivalents in a dose-response study. Shaking responses increased significantly with the dose of either queen-(Fig 5A) or king extracts (Fig 6A), with both 1 and 3 QE treatments and the 1 KE treatment being significantly higher than the respective worker extract controls. Antennation responses to introduced glass dummies were uninformative in these assays. Although termites responded to queen-extracts in a dose-dependent manner (Fig 5B), their antennation responses to 3 QE and worker extracts were not significantly different. Antennation responses to king extracts on glass dummies were not significantly different across all treatments (Fig 6B).

**Fig 5.**
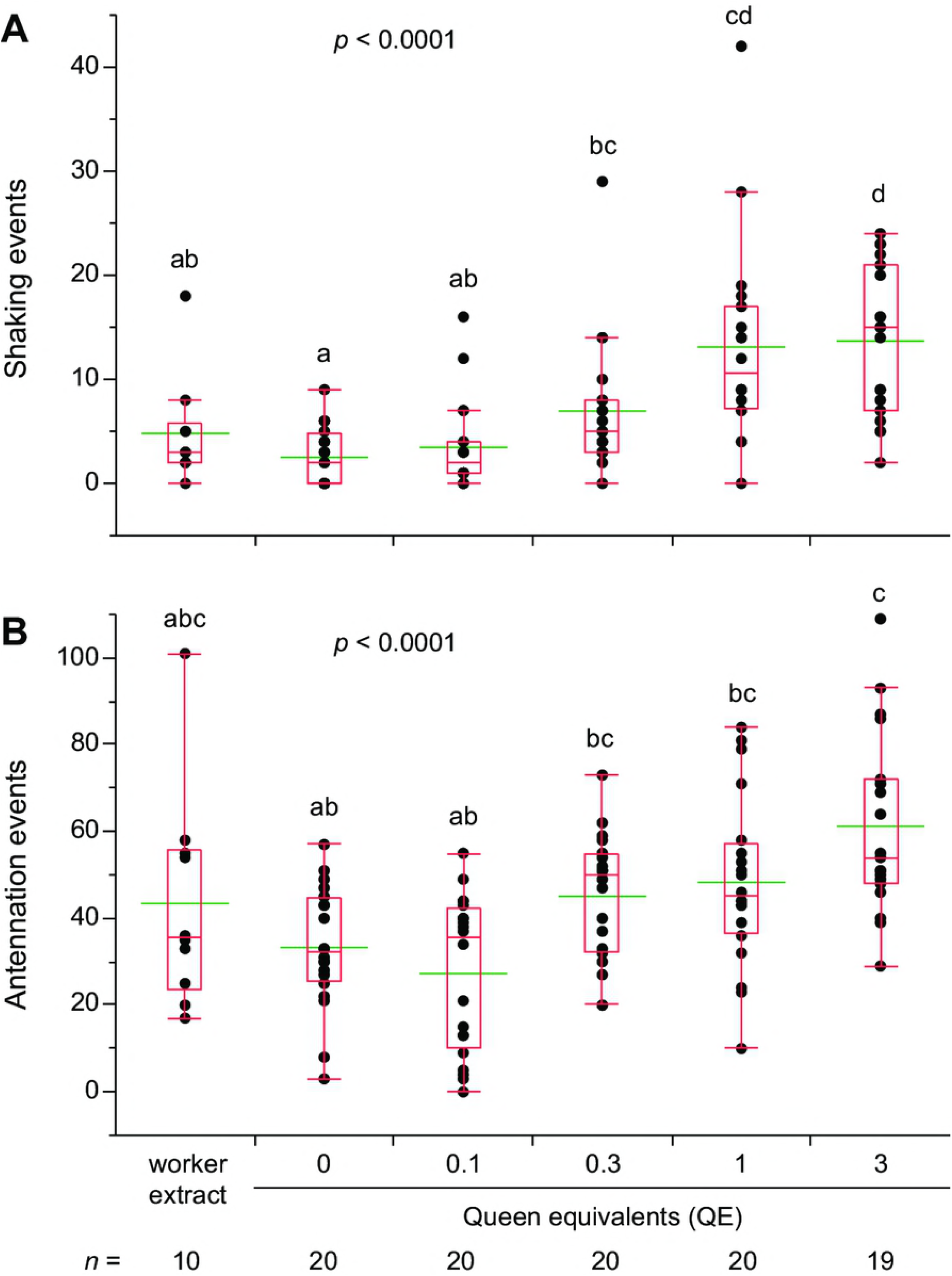
Termite responses to glass dummies treated with hexane extracts of neotenic queens. Lateral shaking (**A**) and antennation (**B**) were measured during 5 min assays for each treatment. Glass dummies were coated with hexane only (0), 0.1, 0.3, 1, and 3 queen equivalents along with worker extracts dissolved in hexane. Hexane extracts of workers were created by pooling 6 workers with mass approximately equal to 3 neotenic queens to approximate the highest queen concentration. Each assay dish consisted of 30 workers, 2 soldiers, and an introduced glass dummy, and assays were conducted under ambient light conditions during the photophase. Letters indicate significantly different values using one-way ANOVA (*p* < 0.05) and Tukey’s HSD. For all treatments the number of replicate assays is indicated under its axis label. In the box plots, the horizontal line within the box represents the median value, the box represents the 25th to 75th quantiles, and the wider green line represents the mean.

**Fig 6.**
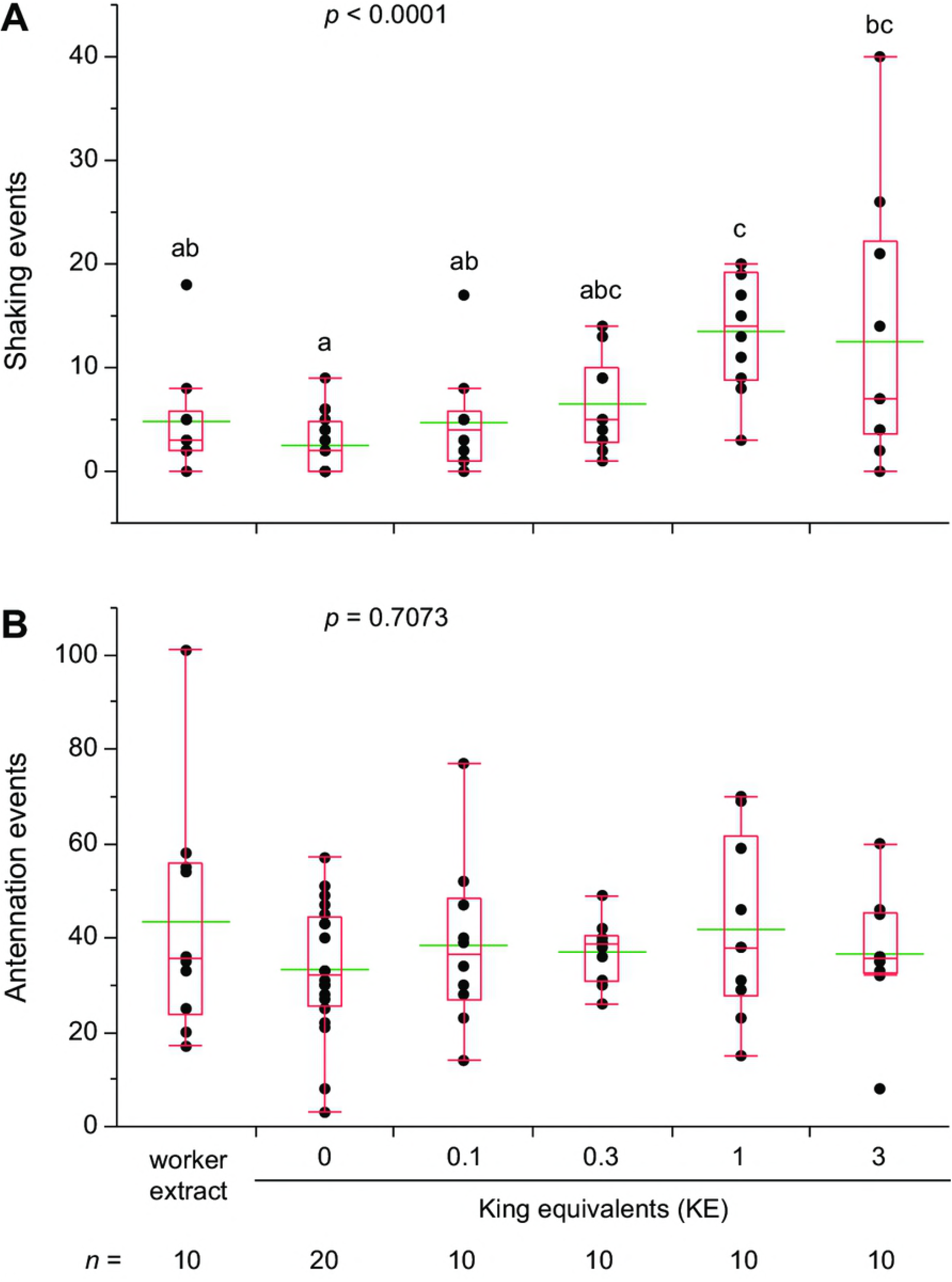
Termite response to glass dummies treated with hexane extracts of neotenic kings. Lateral shaking (**A**) and antennation (**B**) were measured during 5 min assays for each treatment. Glass dummies were coated with hexane only (0), 0.1, 0.3, 1, and 3 king equivalents along with worker extracts dissolved in hexane. Hexane extracts of workers were created by pooling 6 workers with mass approximately equal to 3 neotenic kings to approximate the highest king concentration. Each assay dish consisted of 30 workers, 2 soldiers, and an introduced glass dummy, and assays were conducted under ambient light conditions during the photophase. Letters indicate significantly different values using one-way ANOVA (*p* < 0.05) and Tukey’s HSD. For all treatments the number of replicate assays is indicated under its axis label. In the box plots, the horizontal line within the box represents the median value, the box represents the 25th to 75th quantiles, and the wider green line represents the mean.

The presence or absence of aggression (lunging behavior toward glass dummies) was also recorded in all assays (Fig 7). More aggression (65%) was directed at the control dummies coated with hexane than at workers (20%), kings (0–30% across concentrations), and queens (5–25% across concentrations). All extracts elicited significantly less worker aggression than the control dummies (Chi-square test, workers: df = 1 *p* < 0.017, kings: df = 5, *p* < 0.0001, queens: df = 5, *p* < 0.0001) (Fig 7).

**Fig 7.**
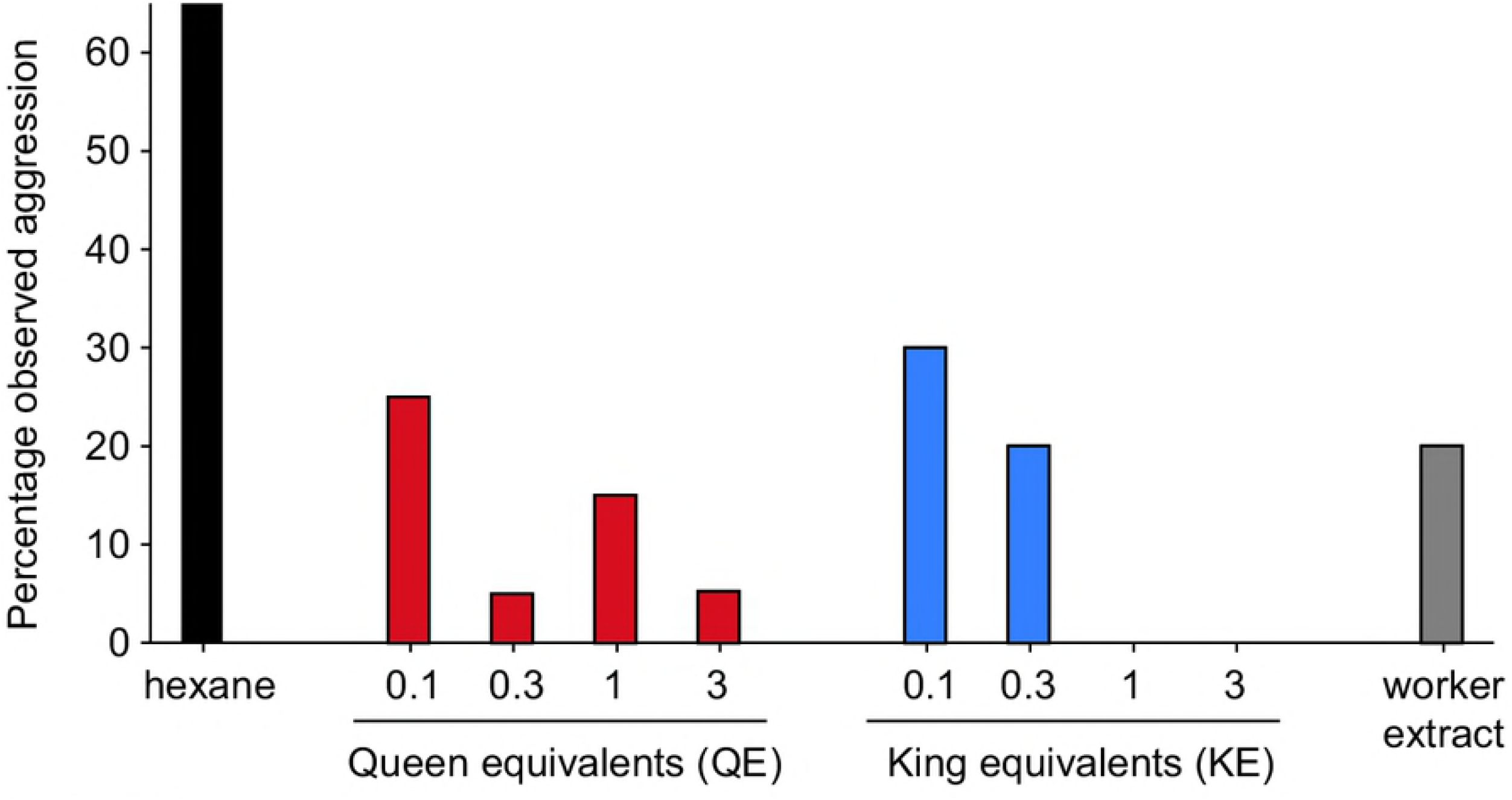
Relationship between aggressive behavior and cuticular extract concentrations of queens, kings, and workers. Hexane was applied to glass dummies for the solvent control treatment. Concentrations are denoted in queen- and king-equivalents applied to glass dummies. Hexane extracts of workers were created by pooling 6 workers with mass approximately equal to 3 queens or kings. Glass dummies (*n* = 20 for all queen concentrations and hexane controls except 3 QEs, which had 19, *n* = 10 for all king concentrations and the worker extract) were observed for 5 min. Chi-square tests for each caste show a significant effect on worker aggression for queens (df = 5, *p* < 0.0001), kings (df = 5, *p* < 0.0001), and workers (df = 1, *p* < 0.017).

## Discussion

### Lateral shaking as a royal recognition behavior

Our aims were to demonstrate the existence of observable and repeatable recognition behavior by workers towards queens and kings in *R. flavipes* and to develop a reliable bioassay to facilitate future isolation and identification of royal recognition pheromones. Our bioassay results strongly support the conclusion that lateral or longitudinal shaking behavior is a strong indicator of neotenic queen and king recognition. Lateral shaking is different from head-drumming, which is used primarily by soldiers either to recruit termites or to send alarm signals through the nest substrate [35,37,39,40]. The lateral shaking behavior was differentially elicited by queens and kings more than by workers in all of our assays. This is the first empirical evidence of behavioral royal recognition in termites, and we used this assay to identify the first royal recognition pheromone in termites [13].

The function of the lateral shaking behavior remains unclear because it occurs in several contexts. First, this behavior is performed away from reproductives and by all castes, including neotenic queens and kings. Secondly, the prevalence of lateral shaking behavior is highly correlated with alarm or disturbance in the colony [41]. Our dark/light assays suggest that this response is intensified under lit conditions and all recognition assays were performed in a lit lab and involved some disturbance as the focal termite or glass dummy was added to the assay dish. Shaking behavior might communicate a rapid local mechanical signal in disturbed or excited conditions to ensure the safety of high-value reproductives or begin repair of damaged areas of the nest. In the drywood termite *Cryptotermes secundus*, workers and nymphs exhibit increased aggression among nestmates after disturbance and an increase in shaking behavior in food-limited situations [31,42]. In both of these cases, the shaking behavior is interpreted as aggressive and it signals a transition from a cooperative to a more self-serving disposition in the study termites.

Shaking most likely does not elicit aggressive behavior in *R. flavipes* in the context of our bioassay. Indeed, aggression in this termite species is less frequent in general than in *Cryptotermes* termites, as contests for replacement reproductives are not commonly observed, the colony is much larger, and the nest habitat is larger and more prone to disturbance in colonies with satellite nests and vulnerable areas outside a single piece of wood. Overall, higher rates of shaking directed toward royals in all our assays, and also toward primary and neotenic queens and kings in undisturbed dishes (CF, personal observation), strongly support the notion that while shaking may serve multiple functions in *R. flavipes*, it is a major and predictable queen and king recognition response. Most significant was the observation that shaking responses increased with the dose of royal extract, whereas aggression responses declined. These results suggest that shaking in the context of this bioassay is a response to royal semiochemicals and not an aggressive response. In other contexts (e.g., foreign workers or soldiers, interspecific interactions) shaking behavior might elicit aggression, but in these situations the shaking response should increase with the dose of the intruder semiochemicals. Because shaking behavior may convey different information in different contexts, it is also plausible that it was co-opted from ancestral alarm or agitation responses that elicited aggressive behaviors to be a royal-recognition response that modulates colony-wide behavior.

Although shaking behavior likely conveys information over relatively short range, as it is typically elicited from physical contact with a reproductive, workers are often observed shaking repeatedly after they move away from the queen or king. Therefore, this behavior could be amplified and dispersed over a longer distance through a chain of workers.

It is also possible that shaking responses in *R. flavipes* vary in response to different stimuli. Our real-time visual observations could not resolve nuances in this behavior, but it is possible that the frequency, amplitude and other features of the behavior may be context-specific. Physical measurements of termite jerking or drumming behavior have been recorded before with few conclusive statements about their purpose [33,35,36]. Whitman and Forschler [37] described four general types of shaking behavior distinguished by speed and frequency in *R. flavipes*. Our assays did not differentiate among these movements but included three of the four described.

Honey bees exhibit a behavior similar to termite shaking, called the vibration response, where individuals shake rapidly, leading their nestmates to change tasks within the colony [43,44]. Bees that receive these vibration stimuli are typically less active and show increased task performance after receiving the signal. Other royal recognition responses in social insects are typically chemically mediated and include retinue responses or other aggregations around royal castes [15,28–30], queen tending behaviors such as grooming or feeding, and strong aggressive responses to establish reproductive dominance or prevent unwanted reproduction in the colony [45,46].

### Royal recognition is chemically mediated in *R. flavipes*

Lateral shaking is readily elicited in *R. flavipes* by cuticular chemicals of neotenic queens and kings (Figs 4–6). We transferred cuticular compounds from queens to worker termites by tumbling them in various queen: worker ratios. We also transferred hexane extracts of queen and king cuticular lipids to glass dummies. In both experiments, the royal-perfumed workers and glass dummies elicited significantly more shaking responses than the respective controls, indicating that royal-recognition pheromones were contained in the transferred chemicals. CHCs are most likely responsible, as they are the dominant feature of insect cuticular lipids, but fatty acids, esters, waxes, or other lipids may be involved. Indeed, in our recent research [13], we identified a suite of CHCs that are highly enriched in *R. flavipes* queens and kings as well as a royal-specific hydrocarbon, *n*-heneicosane. In addition, aggressive behaviors were significantly associated with hexane controls and low concentrations of termite extracts, but not with higher concentrations of queen, king or worker extracts (Fig 7), suggesting that these extracts likely contain colony recognition cues and can mitigate aggressive behaviors toward foreign objects. The behavioral assays we developed and validated for this study also facilitated further experiments that confirmed the activity of one of the candidate royal compounds, *n*-heneicosane, as a recognition pheromone in this species [13].

Other species of termites possess CHCs that have been linked to reproductive status, but their functions in royal recognition have not been demonstrated [12,25]. In contrast, CHC recognition pheromones have been demonstrated in many social hymenopterans, including various ant species and *Polistes* wasps [16,18,28,45,47–49]. Van Oystaeyen et al. [17] found that species from across the hymenopteran phylogeny (ant, bee, and wasp) use similar CHCs as queen pheromones, which act to reduce or suppress ovary development. They also compared fertility signals across 64 species of social Hymenoptera to conclude that saturated CHCs are a conserved class of pheromones that function similarly across a diverse assemblage of species (but see Amsalem et al. [50], countered by Holman et al. [51]). The wide phylogenetic distance between the eusocial Hymenoptera and termites, and their shared use of CHCs as fertility signals, could indicate an intriguing case of convergent evolution that would push the use of CHCs as royal pheromones from ~100 million years ago (evolution of bees, ants and wasps) to ~150 million years ago, when eusocial termites evolved from within the cockroaches.

In conclusion, we report a highly discriminating bioassay that quantitatively related shaking behavior in workers and soldiers to presence of a neotenic queen or king. We further showed that queen and king cuticular compounds elicited this behavior. Our bioassay should prove to be useful for future research to identify specific royal pheromones, the social status of newly emerging reproductives, and the activity of candidate volatile and non-volatile royal pheromones. Queens and kings possess similar cuticular profiles in *R. flavipes* and both sexes elicit increased lateral shaking and antennation. By examining caste-specific differences in cuticular profiles, and using this behavioral assay, we recently identified the chemical basis for this behavior, the first queen recognition pheromone, and the first ever king pheromone in termites [13]. Other caste-specific CHCs remain to be evaluated with these new behavioral assays. Finally, the function of shaking behavior should also be the target of future research to understand how this behavior changes in different contexts within the colony and whether the shaking behavior consists of different elements that require closer scrutiny with high speed photography and laser Doppler vibrometry.

## Supporting Information

**S1 Dataset. Raw assay data for all figures.**

## Acknowledgments

We would like to acknowledge Paul Labadie for help collecting termites. We also would like to thank the administration of Historic Yates Mill County Park, Lake Johnson Park, and Schenck Forest for their support during our project.

